# Local temporal Rac1-GTP nadirs and peaks restrict cell protrusions and retractions

**DOI:** 10.1101/2021.06.23.449555

**Authors:** Jianjiang Hu, Xiaowei Gong, Staffan Strömblad

## Abstract

Spatiotemporal coordination of the GTP-binding activity of Rac1 and RhoA initiates and reinforces cell membrane protrusions and retractions during cell migration^1–7^. However, while protrusions and retractions form cycles that cells use to efficiently probe their microenvironment^8–10^, the control of their finite lifetime remains unclear. To examine if Rac1 or RhoA may also control protrusion and retraction lifetimes, we here define the relation of their spatiotemporal GTP-binding levels to key protrusion and retraction events, as well as to cell-ECM mechanical forces in fibrosarcoma cells grown on collagen of physiologically relevant stiffness. We identified temporal Rac1-GTP nadirs and peaks at the maximal edge velocity of local membrane protrusions and retractions, respectively, followed by declined edge velocity. Moreover, increased local Rac1-GTP consistently preceded increased cell-ECM traction force. This suggests that Rac1-GTP nadirs and peaks may restrain the lifetime of protrusions and retractions, possibly involving the regulation of local traction forces. Functional testing by optogenetics validated this notion, since local Rac1-GTP elevation applied early in the process prolonged protrusions and restrained retractions, while local Rac1-GTP inhibition acted in reverse. Optogenetics also defined Rac1-GTP as a promotor of local traction force. Together, we show that Rac1 plays a fundamental role in restricting the size and durability of protrusions and retractions, plausibly in part through controlling traction forces.

## Main

Cell membrane protrusions and retractions are highly dynamic membrane structures involved in cell-extracellular matrix (ECM) interactions, cell migration and invasion^4, 11, 12^. Constant membrane protrusion and retraction formation in different positions of the cell leads to directed cell migration, while local dynamic cell membrane protrusion-retraction cycles provide cells with probing, exploratory ability of the surrounding microenvironment^8–10^. The Rho family small GTPases RhoA and Rac1 cycles between GTP-bound and GDP-bound states and are central regulators of cytoskeletal dynamics, cell-ECM adhesions and myosin contractions^6, 7, 13, 14^. Coordination of their GTP-bound activities governs protrusion and retraction initiation and reinforcement. Rac1-GTP promotes local cell-ECM adhesion and actin filament assembly that drive cellular protrusions^7, 13, 15^, while RhoA-GTP promotes cell-ECM adhesion maturation and stimulates interactions between myosin and actin filaments to drive local contractility. This way, RhoA promotes the initiation and reinforcement of both protrusions and retractions^16–18^. Thus, Rac1-GTP and RhoA-GTP produce cell shape changes with force application at cell-ECM adhesions^16, 19–21^. Local protrusive and contractile forces in turn redistribute or reorganize actin filaments, adhesions, and their regulators^22–24^. However, it remains unclear how initiated and reinforced protrusions and retractions may be restrained to facilitate the naturally occurring dynamic cycles of local protrusions and retractions and if this restrain might involve local alterations of cell-ECM forces.

The mechanics of the cellular microenvironment, including the substrate stiffness affects cell membrane behavior and signaling^23–26^. Because conventional glass/plastic surfaces are in the million-fold range stiffer (1-7 GPa) than mammalian tissues^27^, we provided HT1080 human fibrosarcoma cells with collagen type I of physiologically relevant stiffness coated onto a polyacrylamide (PAA) gel (6.9 kPa)^28, 29^ also surface-labeled with red fluorescent beads to accommodate traction force measurements^30, 31^ (Fig. 1a). Genetically encoded FRET-based biosensors were utilized to capture Rac1-GTP or RhoA-GTP levels as well as the membrane protrusion and retraction dynamics ^19, 32, 33^. 3D FRET imaging with lattice light sheet microscopy revealed that the vast majority of the Rac1-GTP and RhoA-GTP located at the cell-ECM interface (Supplementary Fig. 1, 2). High RhoA-GTP was also observed at membrane ruffles (Supplementary Fig. 3), as previously suggested^19^.We then used conventional confocal microscopy to simultaneously capture Rho GTPase activities and traction forces at sub-micron resolution with 10 s time interval near the cell-ECM interface.

**Figure 1.**
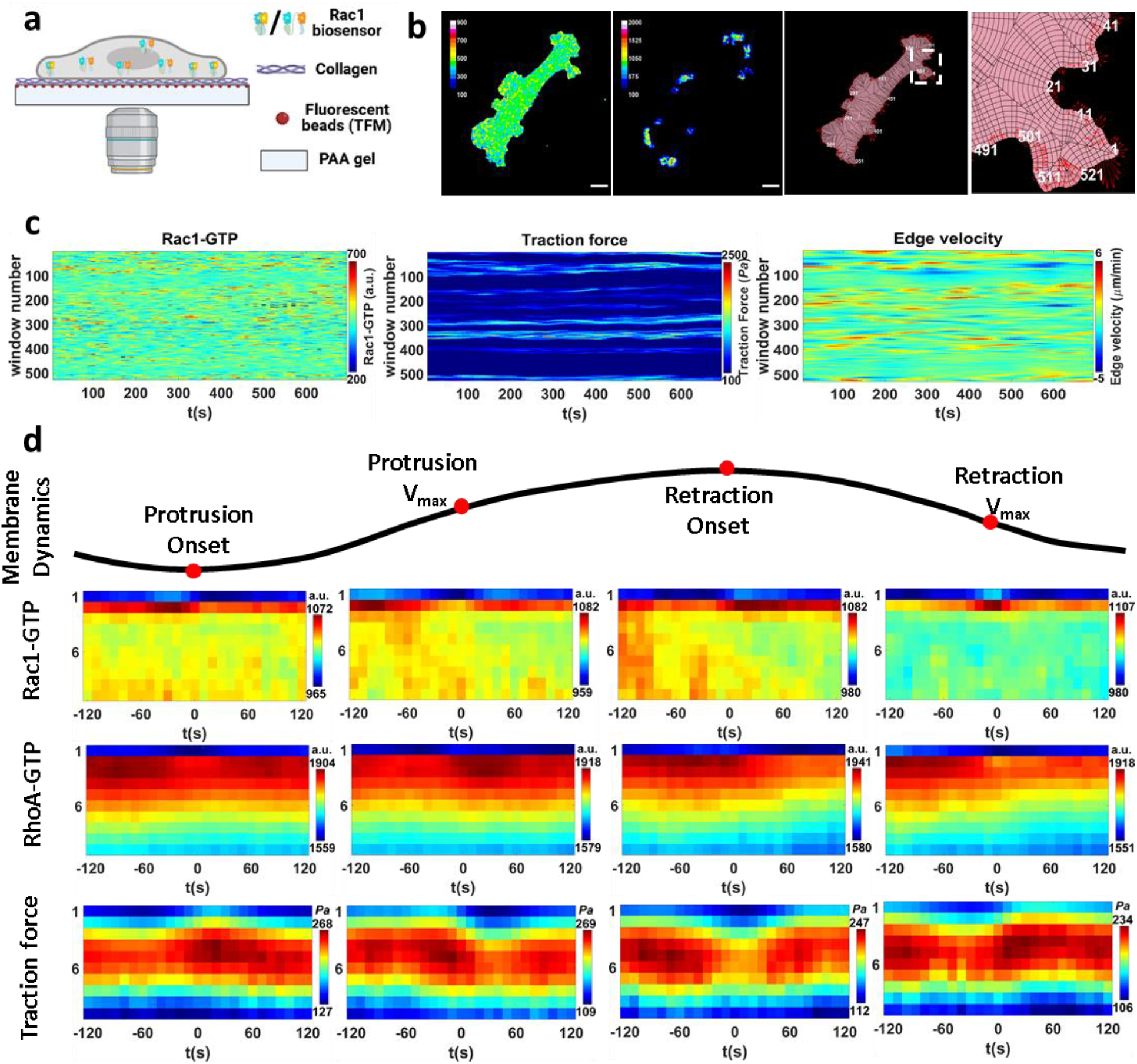
Spatiotemporal analysis of Rac1-GTP, RhoA-GTP and traction forces at key cell membrane events. **a-c. High resolution imaging and analysis simultaneously capturing GTP-bound activity, traction force and cell membrane movements.** a) Schematic image of the platform for simultaneous measurement of highly resolved GTP-binding activity, cell traction force and cell membrane movements. HT1080 human fibrosarcoma cells stably expressing genetically encoded FRET biosensors were utilized to capture Rac1-GTP or RhoA-GTP levels. Cells were seeded onto a Collagen type I coated surface with a physiologically relevant stiffness (Young’s modulus 6.9 kPa) also surface-labeled with red fluorescent beads to accommodate traction-force measurements. Signals were captured via the objective at the bottom providing sub-micron resolution. b) The sample images show the acquired Rac1-GTP level (left), traction force (middle left), outlay of the window sampling (middle right) and zoom-in of the window sampling (right). Rac1-GTP (a.u.) and traction force (*Pa*) levels are shown in pseudo colors according to the inset scales. For the window sampling images, the pink area shows the segmented cell area and the black lines show how the cells are sampled into 1 μm wide cell edge windows based on local cell edge geometry. Windows are numbered in white along the cell edge around the cell. Red arrows show the instant edge sector velocity indicating cell membrane movement direction and velocity (length of arrows). Scale bar 20 μm. c) GTPase activity, traction force and cell edge velocity. Registered and smoothed temporal dynamics of GTP-bound activity (left) and traction force (middle) levels in the second windows (1-2 μm from the cell edge) all around the cell displayed over time as kymographs, with the corresponding edge velocity (right). All graphs are shown in pseudo color according to the indicated scales. **d. Rac1-GTP, RhoA-GTP and traction force levels around key cell membrane events.** The dynamics of segmented cell membrane edge sectors (1 μm wide) were aligned according to cell membrane Protrusion onset, maximal protrusion velocity (Protrusion V_max_), Retraction onset, and maximal retraction velocity (Retraction V_max_). The mean values of Rac1-GTP, RhoA-GTP and traction force levels at different depths of the cell (1-10 μm, y axis) around the specific time points (−120 s to + 120 s, x axis) are shown in pseudo color. Sample size: Rac1-GTP & traction force: 5852 protrusions and 3817 retractions from 13 cells; RhoA-GTP: 8732 protrusions and 4244 retractions from 23 cells.

To analyze our time-lapse images, we applied the window sampling method developed by the Danuser lab^5^ to segment the entire cell into 1 μm × 1 μm windows that were tracked over time. This derived time series of local membrane edge dynamics as well as the corresponding Rac1-GTP/RhoA-GTP and traction force levels in each window from the cell edge to the cell center (Fig. 1b, c and Supplementary Mov. 1). By aligning and averaging thousands of time series, we obtained a quantitative measure of the relation between the highly localized protrusion/retraction membrane dynamics and the dynamic alterations of Rac1-GTP, RhoA-GTP and traction force levels.

We aligned the time series of all windows from the different cells according to four different edge motion events^5^: protrusion onset, maximal protrusion velocity (protrusion V_max_), retraction onset and maximal retraction velocity (retraction V_max_). This allowed us to quantify the dynamics of the mean Rac1-GTP, RhoA-GTP and traction force levels at different depth layers of the cell at the time periods surrounding each of these four edge motion events (Fig. 1d). Switching the sector width from 1 μm to 5 μm did not affect the patterns of the result maps, displaying robustness of the results over different window sizes (Supplementary Fig. 5). We found that the majority of Rac1-GTP and RhoA-GTP signals located within 2 μm from the cell edge, while the highest traction force level located 4-6 μm away from the cell edge, with a lower but correlating traction force at 1-2 μm depth (Supplementary Fig. 6). The universally low-level signals in the first window (0-1 μm away from the segmented cell edge) was most likely caused by segmentation imperfection. To avoid edge segmentation errors to cause artificial data fluctuations while entailing the most meaningful signals, we used the Rac1-GTP, RhoA-GTP and traction force signals from the second sampling windows (1-2 μm away from the segmented cell edge) for further analysis (Fig. 2a-e).

**Figure 2.**
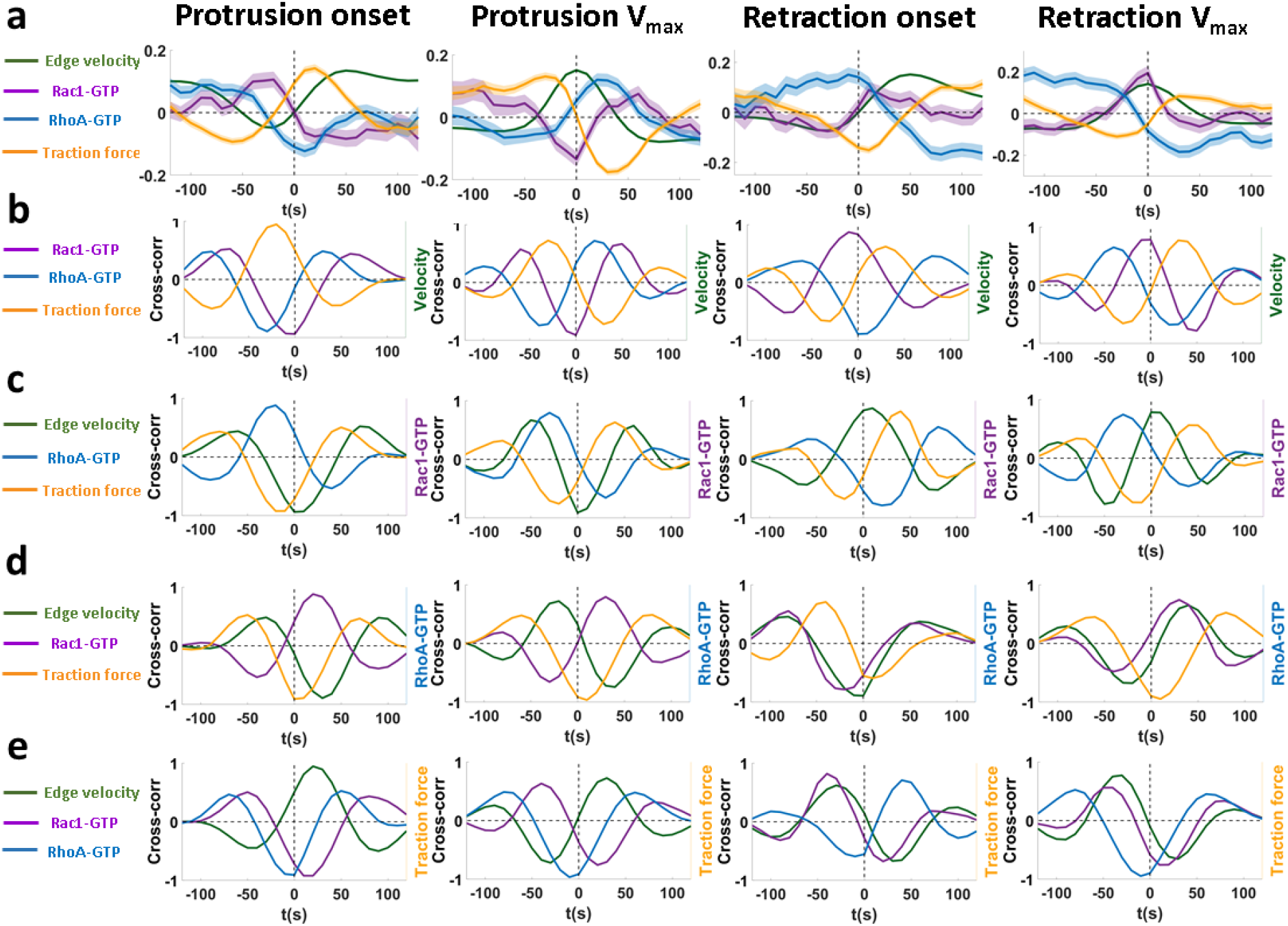
Localized temporal Rac1-GTP nadirs and peaks at the cell membrane maximum velocity of protrusions and retractions and their correlations with RhoA-GTP and traction forces. **a. Rac1-GTP, RhoA-GTP and traction force levels around key cell membrane events.** The mean values of Rac1-GTP, RhoA-GTP and traction force levels at the second window (1-2 μm from the segmented cell edge) before and after the Protrusion onset, Protrusion V_max_, Retraction onset and Retraction V_max_, extracted from the data represented in Figure 1d. The raw data were normalized with the z-score method before calculating the mean values. Solid lines show the mean values and the shadows show the 95% confidence intervals. Sample size: Rac1-GTP, velocity & traction force: 5852 protrusions and 3817 retractions from 13 cells; RhoA-GTP: 8732 protrusions and 4244 retractions from 23 cells. **b-e. Cross-correlation analyses between cell edge velocity, Rac1-GTP, RhoA-GTP and Traction force.** Cross-correlation analysis of mean Rac1-GTP (purple), RhoA-GTP (blue), traction force (orange) levels in the second window (1-2 μm from the segmented cell edge), and corresponding cell edge sector mean velocity (green) towards each other around Protrusion onset, Protrusion V_max_, Retraction onset and Retraction V_max_. The cross-correlation analysis is centered on (b) Cell edge velocity; (c) Rac1-GTP levels; (d) RhoA-GTP levels and (e) traction force levels. Cross-correlation results are shown in colored lines as indicated on the left side of each graph.

At the protrusion onset, high RhoA-GTP levels at the cell edge arose 20-30 s before an elevation of Rac1-GTP levels, in turn occurring approximately 40 s before the protrusion onset that was paralleled by increased local traction force (Fig. 2a). High RhoA-GTP also appeared prior to the retraction onset, while traction force levels decreased before the retraction onset (Fig. 2a). These Rac1 and RhoA dynamics are consistent with previous studies of protrusion and retraction initiation performed on glass/plastic surfaces with approximately million-fold higher stiffness^3, 19, 32^, suggesting robustness of cell membrane dynamics regulation across environments with different mechanical properties.

Importantly, Rac1-GTP levels gradually decreased after protrusion onset to reach a nadir at the protrusion V_max_ (Fig. 2a). Thus, a Rac1-GTP nadir at the protrusion V_max_ occurs at a restriction point, after which the membrane velocity slows down. Conversely, Rac1-GTP increased during retractions to reach a peak at the retraction V_max_ (Fig. 2a), suggesting that a Rac1-GTP peak occurs at the restriction point for cell membrane retractions.

Interestingly, Rac1-GTP levels displayed a positive correlation with traction force levels around each of these four membrane movement events with Rac1-GTP alterations preceding traction force changes by approximately 40 s (Fig. 2a, e). Rac1-GTP level alterations were also followed by RhoA-GTP level changes in the opposite direction (Fig. 2a, d). Unexpectedly and contrary to Rac1-GTP, the local RhoA-GTP level fluctuations displayed a negative correlation with local traction force alterations around each of the four membrane movement events (Fig 2a, e).

The local Rac1-GTP nadirs and peaks at the protrusion and retraction V_max_ made us hypothesize that Rac1-GTP levels may play a central role in restraining membrane protrusions and retractions. We therefore postulated predictions based on this hypothesis that could be used for functional testing by optogenetics based local Rac1-GTP perturbations. We predicted how local induction of Rac1-GTP or Rac1-GDP (inhibiting Rac1-GTP) before protrusion or retraction V_max_ would affect the velocity and the duration of protrusions and retractions, as well as its influence on local traction force (Fig. 3a). There is also the alternative possibility that the Rac1-GTP level changes are the consequence of the membrane dynamics, rather than executing a regulatory role. This alternative would predict no membrane dynamic changes upon Rac1-GTP perturbation. According to our predictions in Fig 3a, we designed and performed Rac1 perturbation experiments with genetically encoded optogenetic tools. Photoactivatable “constitutively active” (GTP-bound) Rac1 (PA-Rac1(Q61L)) or “dominant negative” (GDP-bound) Rac1 (PA-Rac1(T17N))^34^ were induced by blue light in a small region of the HT1080 cells (Fig. 3b). As predicted by our hypothesis, activation of PA-Rac1(Q61L) enhanced membrane protrusions and restricted retractions, while activation of PA-Rac1(T17N) restricted the ongoing protrusions and enhanced the retractions (Fig. 3c, d).

**Figure 3.**
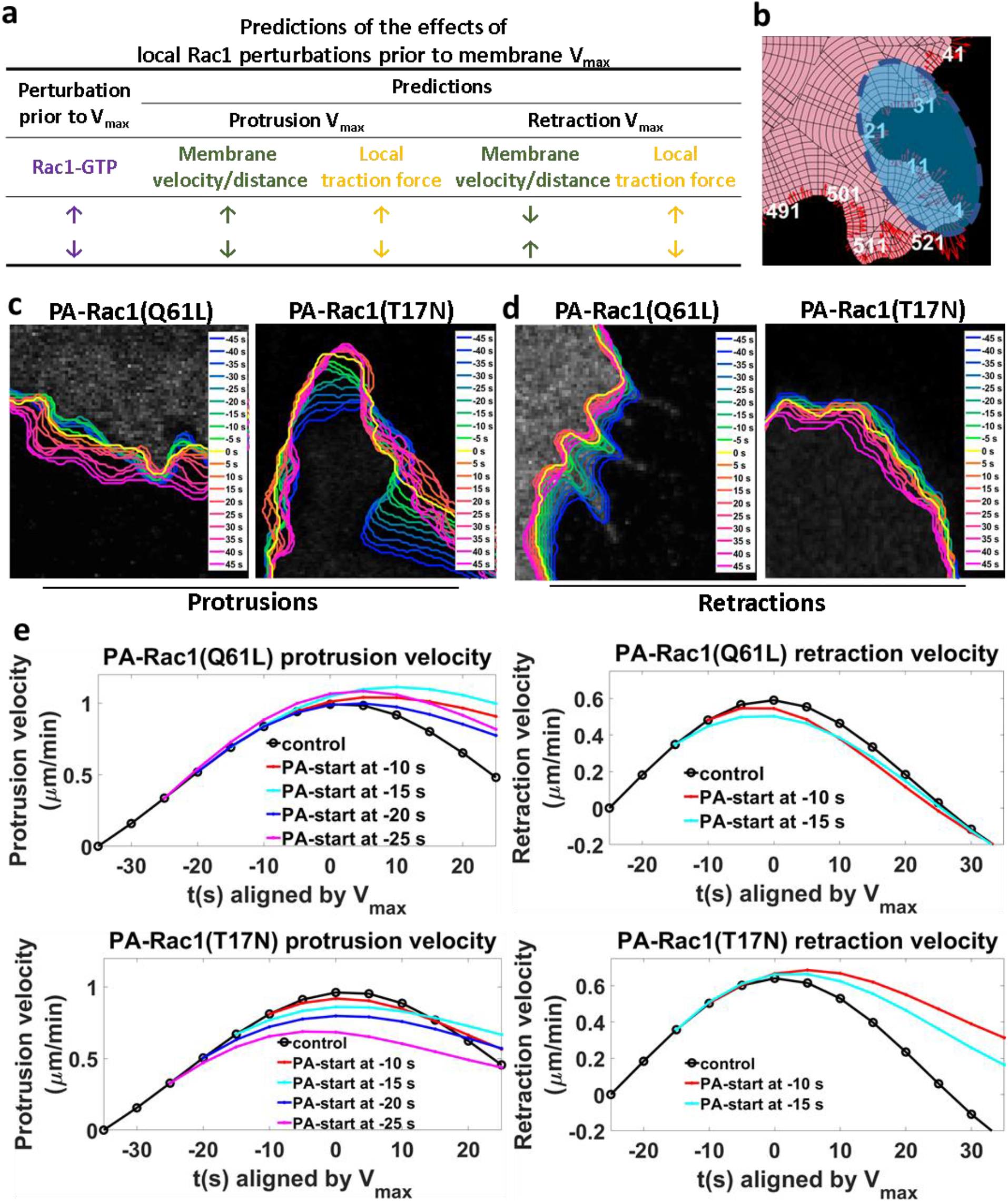
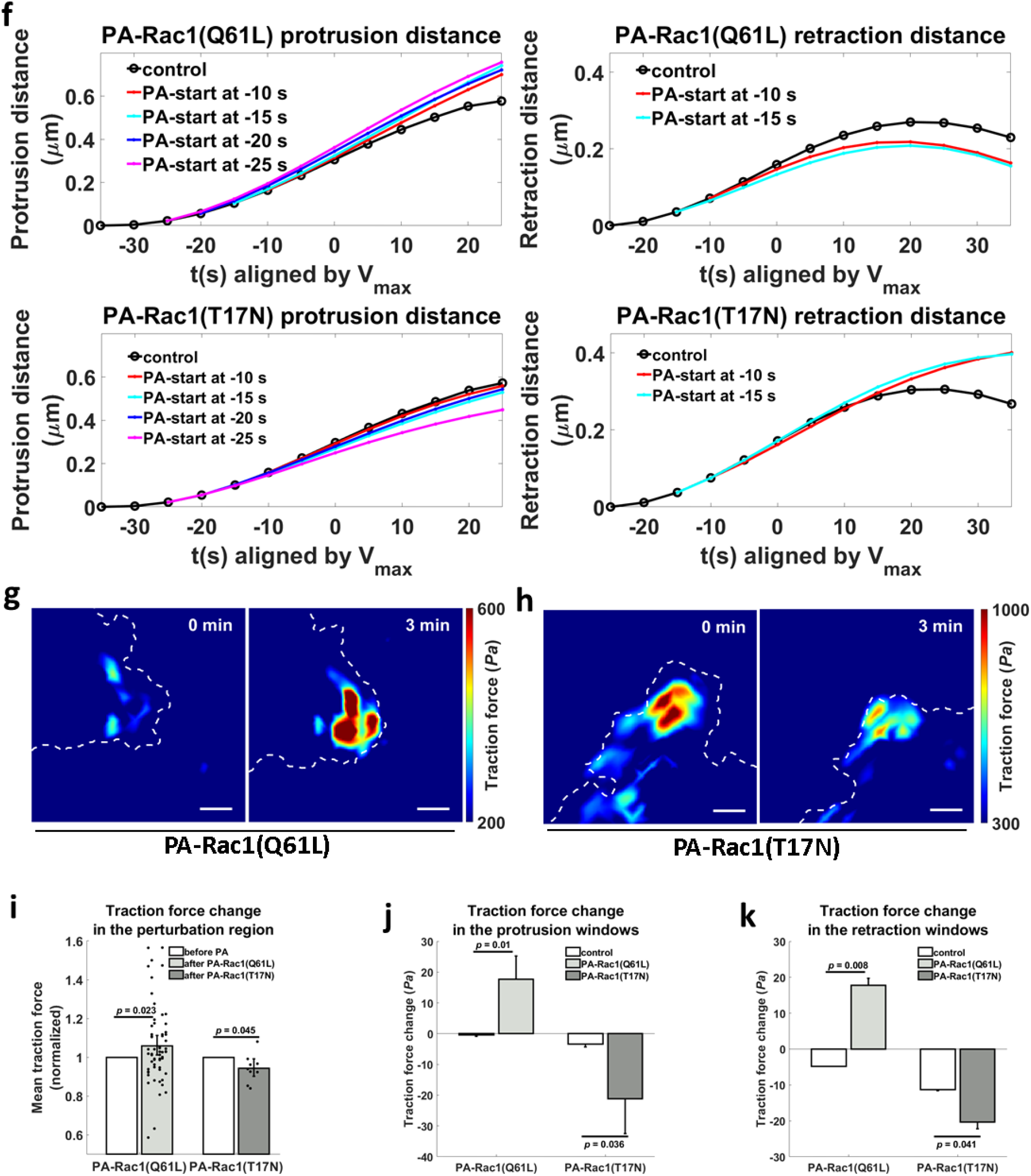
Functional role for Rac1 in the restriction of protrusions and retractions. **a. Predictions of the effects of local Rac1 perturbations on cell membrane protrusions and retractions.** Predictions on the cell membrane movements are based on the hypothesis that the Rac1-GTP nadirs and peaks that we observed at Protrusion V_max_ and Retraction V_max_ functionally restrict protrusions and retractions. The table indicates how activation (purple upwards arrows) or inhibition (purple downwards arrows) of Rac1-GTP before V_max_ would alter cell membrane velocity V_max_ and total membrane movement distance (green arrows), as well as the local traction force (yellow arrows). Arrow directions indicate predicted directions of change. Summary of the predictions: Local increase in Rac1-GTP levels prior to protrusion V_max_ will lead to later and higher protrusion V_max_ and longer protrusion distance. Inhibition of Rac1-GTP prior to protrusion V_max_ will lead to earlier and lower protrusion V_max_ and shorter protrusion distance. Increase in Rac1-GTP levels prior to retraction V_max_ will lead to earlier and lower retraction V_max_ and shorter retraction distance. Inhibition of Rac1-GTP levels prior to retraction V_max_ will lead to later and larger retraction V_max_ and longer retraction distance. Local increase in Rac1-GTP levels will increase the local traction force level in both protrusions and retractions, while inhibition of Rac1-GTP will decrease the local traction force levels in both protrusions and retractions. **b-k. Experimental validation of predictions by local optogenetic Rac1 perturbations.** Local Rac1-GTP activity in a small region of the cells was elevated (PA-Rac1(Q61L)) for 1 min or inhibited (PA-Rac1(T17N)) for 2 min with blue light. Cell membrane dynamics and traction force were imaged and quantified by confocal microscopy at 5 s intervals before, during and after the perturbations. **b. Schematic image of optogenetic Rac1 perturbation.** The local blue region was subject to blue laser light to optically induce a Rac1 perturbation. **c-d. Sample time lapse images of membrane protrusions and retractions upon Rac1 perturbations.** Photoactivation of Rac1-GTP (PA-Rac1(Q61L)) or Rac1-GDP (PA-Rac1(T17N)) was performed in protruding (c) or retracting (d) cell areas. Cell images 45 s before photoactivation start shown in gray and the following cell edge dynamics in pseudo-colors coding for different time points as indicated. Yellow shows the photoactivation starting time point. Cold and warm colors show the cell edge location before and after the start of the photoactivation at the time points indicated in the insets. **e-f. Local Rac1-GTP levels restrict protrusions and retractions.** Rac1-GTP (PA-Rac1(Q61L)) or Rac1-GDP (PA-Rac1(T17N); inhibiting Rac1-GTP)) was photoactivated at indicated times before the cell membrane V_max_ of protrusions and retractions. Mean values of membrane protrusion and retraction velocities (e) and distances (f) before and after V_max_ are plotted with the difference in difference (DID) method. Black lines (control) show the mean protrusion or retraction velocities/distances of the membrane edge sectors in the perturbation region before (without) photoactivation. Colored lines show the difference of mean membrane velocities/distances towards controls upon photoactivation. **(Left)** Perturbations started 10 s (red), 15 s (cyan), 20 s (blue) or 25 s (magenta) before the theoretical Protrusion V_max_ time point. Statistical analysis using the false discovery rate (FDR) method^44, 45^ showed that, except for the - 10 s PA-Rac1(T17N) membrane protrusion distance, the Q values of all the other perturbations are smaller than the 0.1% threshold when compared to the corresponding controls. Sample details : *PA-Rac1(Q61L):* control (n = 1545); −10 s (n = 97, velocity (Q = 5.05×10^−3^, 0.1% threshold = 9.99×10^−1^), distance (Q = 9.95×10^−1^, 0.1% threshold = 9.99×10^−1^)); −15 s (n = 86, velocity (Q = 1.38×10^−3^, 0.1% threshold = 9.99×10^−1^), distance (Q = 9.92×10^−1^, 0.1% threshold = 9.99×10^−1^)); −20 s (n = 82, velocity (Q = 9.81×10^−1^, 0.1% threshold = 9.99×10^−1^), distance (Q = 9.97×10^−1^, 0.1% threshold = 9.99×10^−1^)); −25 s (n = 75, velocity (Q = 7.11×10^−1^, 0.1% threshold = 9.99×10^−1^), distance (Q = 9.95×10^−1^, 0.1% threshold=9.99×10^−1^)) from 83 cells; *PA-Rac1(T17N):* control (n = 1127); −10 s (n = 60, velocity (Q = 9.96×10^−1^, 0.1% threshold = 9.99×10^−1^), distance (Q = 9.99×10^−1^, 0.1% threshold = 9.99×10^−1^)); −15 s (n = 62, velocity (Q = 9.85×10^−1^, 0.1% threshold = 9.99×10^−1^), distance (Q = 9.997×10^−1^, 0.1% threshold = 9.999×10^−1^)); −20 s (n = 49, velocity (Q = 9.88×10^−1^, 0.1% threshold = 9.99×10^−1^), distance (Q = 9.997×10^−1^, 0.1% threshold = 9.999×10^−1^)); −25 s (n = 73, velocity (Q = 8.33×10^−1^, 0.1% threshold = 9.99×10^−1^), distance(Q = 9.98×10^−1^, 0.1% threshold = 9.99×10^−1^)) from 69 cells. **(Right)**Perturbations started 10 s (red), 15 s (cyan) before the theoretical Retraction V_max_ time point. Statistical analysis using the false discovery rate (FDR) method showed that the Q values of all the perturbations in the graph are smaller than the 0.1% threshold when compared to the corresponding controls. Sample sizes: *PA-Rac1(Q61L):* control (n = 162); −10 s (n = 52, velocity (Q = 9.987×10^−1^, 0.1% threshold = 9.998×10^−1^), distance (Q = 9.998×10^−1^, 0.1% threshold = 9.999×10^−1^)); −15 s (n = 53, velocity (Q = 9.987×10^−1^, 0.1% threshold = 9.998×10^−1^), distance (Q = 9.997×10^−1^, 0.1% threshold = 9.999×10^−1^)) from 83 cells; *PA-Rac1(T17N):* control (n = 120); −10 s (n = 60, velocity (Q = 1.81×10^−3^, 0.1% threshold = 9.99×10^−1^), distance (Q = 9.90×10^−1^, 0.1% threshold = 9.99×10^−1^)); −15 s (n = 59, velocity (Q = 8.78×10^−2^, 0.1% threshold = 9.99×10^−1^), distance (Q = 9.94×10^−1^, 0.1% threshold = 9.99×10^−1^)) from 69 cells. **g-h. Sample images of local traction force upon Rac1-GTP level perturbation.** Local traction forces before and 3 min after photoactivation of Rac1-GTP (PA-Rac1(Q61L), g) or Rac1-GDP (PA-Rac1(T17N), h) are shown in pseudo-colors. Dashed white lines show the cell edge. All of these regions are within the photoactivation area. Scale bar: 5 μm. **i-k. Rac1-GTP levels affects local cell traction force.** (i) Mean traction force in the photoactivation region 3-4 min after the start of induction of PA-Rac1(Q61L) or PA-Rac1(T17N) compared to the mean traction force 0-1 min before induction. *p* values based on paired-sample *t*-test. Results were derived from 60 cells (PA-Rac1(Q61L)) and 9 cells (PA-Rac1(T17N)), respectively. (j-k) Mean traction force change in the second windows (1-2 μm from the segmented cell edge) of the corresponding protrusions (j) or retractions (k) sectors 2 min after the start of induction of PA-Rac1(Q61L) or PA-Rac1(T17N) compared to the mean traction force change in the windows of the sectors before the perturbation started, at time points when these sectors displayed corresponding phases of protrusions or retractions. *p* values based on two-sample *t*-test. PA-Rac1 (Q61L) results were derived from 9178 (control) or 641 (perturbed) protrusions and 925 (control) or 166 (perturbed) retractions of 83 cells. PA-Rac1(T17N) results were derived from 6452 (control) or 385 (perturbed) protrusions and 619 (control) or 188 (perturbed) retractions of 69 cells.

The window sampling method provided detailed quantitative results of the time lapse images, which allowed us to quantitatively compare the local membrane behavior differences before and after the local photoactivation. Induction of PA-Rac1(Q61L) before protrusion V_max_ extended the time until V_max_ was reached and also increased the V_max_ value. Consequently, the membrane protrusion distance was extended. The longer time the PA-Rac1(Q61L) was induced before the membrane protrusion V_max_, the stronger was the protrusion elevation (Fig. 3e, f, left). On the contrary, induction of PA-Rac1(T17N) before protrusion V_max_ led to earlier V_max_ arrival time, lower V_max_, and a shorter protrusion distance. Also in this case, the longer time before protrusion V_max_ that PA-Rac1(T17N) was induced, the earlier and smaller V_max_ and the shorter protrusion distance were observed (Fig. 3e, f, left). Thus, replacing the Rac1-GTP nadir occurring at the protrusion V_max_ with induction of Rac1-GTP caused prolonged protrusions, while enhancing the Rac1-GTP nadir through induction of Rac1-GDP further restricted the protrusions. This strongly indicates that the Rac1-GTP nadir functions to restrict protrusions.

Induction of PA-Rac1(Q61L) before retraction V_max_ shortened the time to V_max_, lowered the V_max_ and shortened the membrane retraction distance, all with a stronger effect the earlier the Rac1-GTP activation was added (Fig. 3e, f, right). On the contrary, local induction of PA-Rac1(T17N) before retraction V_max_ delayed V_max_ occurrence, increased V_max_, and prolonged the retraction distance (Fig. 3e, f, right). This means that enhancement of the Rac1-GTP peak occurring at retraction V_max_ further restricted the retractions, while counteracting this Rac1-GTP peak reversed the naturally occurring restriction. This indicates a regulatory role for the Rac1-GTP peak in the restriction of membrane retractions.

Our finding of a consistent positive cross-correlation where Rac1-GTP preceded traction force around the key membrane events (Fig. 2a, e), suggest that Rac1-GTP may induce traction force. Importantly, this suggestion was functionally validated, since optogenetic activation of PA-Rac1(Q61L) increased traction force levels, while PA-Rac1(T17N) inhibited the traction force within the cellular region exposed to optical PA-Rac1(T17N) activation (Fig. 3g-i). The window sampling based quantification showed that in both protrusion and retraction windows, induction of PA-Rac1(Q61L) increased the local traction force while PA-Rac1(T17N) decreased the local traction force (Fig. 3j, k). Together, this defines a role for Rac1-GTP in the promotion of traction forces.

We here define a broad role for Rac1 in the control of cell membrane dynamics, confirming its role in membrane protrusion initiation^4, 7^ and assigning new functions for Rac1 in the restriction of both membrane protrusions and retractions. Thus, highly localized and temporally precise regulation of Rac1-GTP levels appears to be central for the dynamic membrane protrusion-retraction cycles that cells use to probe the microenvironment^35, 36^. Consistently, Rac appears critical for the probing behavior in *Dictyostelium*^37^.

We conclude that local Rac1-GTP nadirs limit cell membrane protrusions. Low Rac1-GTP levels are linked to sparse non-networked membrane-proximal F-actin^15^ and limited nascent adhesions^13^. Given that both F-actin meshwork density and cell-matrix adhesions provide forces supporting cell membrane protrusions, we propose that the low Rac1-GTP at V_max_ may cause reduced membrane support that restrains the local protrusion velocity (Fig. 4 left). The Rac1-GTP peak-mediated restriction of cell membrane retractions may work in the opposite manner. High Rac1-GTP is linked to a dense membrane-proximal F-actin meshwork^15^ and abundant nascent adhesions^13^. We thereby infer that the high Rac1-GTP at retraction V_max_ may result in abundant F-actin and cell-matrix adhesions which reduce the local retraction velocity by their strong supporting force to resist membrane retraction (Fig. 4 right). However, while Fam49/CYRI- and ARHGAP39-mediated Rac1-inhibition can inhibit protrusions and a large number of GEFs, GAPs and GDIs can control Rac1-GTP levels^6, 13^, it remains to be investigated how Rac1-GTP levels may be regulated to form local temporal nadirs and peaks during the heights of protrusions and retractions.

**Figure 4.**
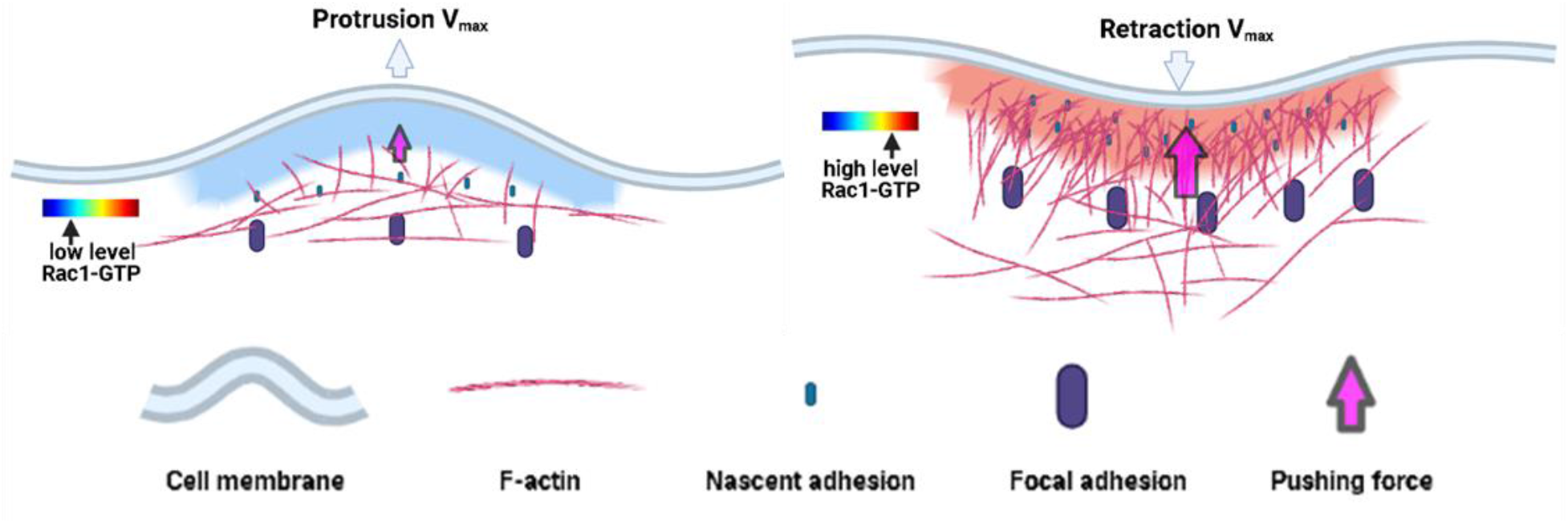
Schematic models for the role of Rac1 in restricting cell protrusions and retractions. Cell membrane protrusions and retractions are large transient structures used for probing of the microenvironment where the cell membrane locally protrudes or retracts usually followed by a reversion. After protrusion/retraction initiation, the local cell membrane velocity gradually increases until it reaches a peak (V_max_) that represents a restriction point, at which the protrusion/retraction is restrained and then reverted. We here found a role for Rac1-GTP activity in restricting membrane protrusions and retractions and that Rac1-GTP promotes local cellular traction forces. **Left:** We found that Rac1-GTP near the cell edge reaches high levels ~20-30 s before protrusion initiation. Then, Rac1-GTP levels gradually decline to reach a nadir at protrusion V_max_, a nadir we found to functionally restrict protrusions. Low Rac1-GTP levels are linked to sparse non-networked membrane-proximal F-actin^15^ and limited nascent adhesions^13^. Given also that both F-actin meshwork density and cell-matrix adhesions provide forces supporting cell membrane protrusions, we hypothesize that the low Rac1-GTP at V_max_ causes reduced membrane support that restrains the local protrusion velocity. **Right:** We found that Rac1-GTP was low at the initiation of retractions and then gradually increased to peak at retraction V_max_, at which the Rac1-GTP peak restricts retractions. High Rac1-GTP is linked to a dense membrane-proximal F-actin meshwork^15^, abundant nascent adhesions^13^ and high cell traction force (Fig. 3g). We therefore hypothesize that the high Rac1-GTP at retraction V_max_ results in abundant F-actin and cell-matrix adhesions that reduce the local retraction velocity by their supporting force to resist membrane retraction.

We report simultaneous FRET biosensor and traction force microscopy (TFM) measurements at sub-micron resolution, facilitated by applying the TFM beads at the surface of the PAA-gel, compared to conventional TFM-beads embedded within the gel that prevents concomitant FRET measurements. This way, we identified a time-lagged positive correlation between Rac1-GTP and traction force levels. By use of optogenetic Rac1 activation, we then defined a functional role of Rac1-GTP to promote traction forces. This Rac1 promotion of traction forces may be brought about by the known function of Rac1 to promote F-actin networks and cell-matrix adhesions, both critical for the generation of traction force^26, 35^. This capability of Rac1 to promote traction force may contribute to the here identified role of Rac1 to restrict protrusions and retractions (Fig 4).

In summary, we found that local Rac1-GTP temporal fluctuations control the local membrane edge velocities that are critical for restricting the size and durability of protrusions and retractions.

## Methods

### Cell line and culture

Human fibrosacoma cell line HT1080 (identified and obtained from Erik Sahai’s lab, Francis Crick Institute) was cultured in DMEM (Gibco) supplemented with 10% fetal bovine serum (FBS, Gibco), 1 mM sodium pyruvate (Gibco), 100 units ml^−1^ penicillin and 100 μg ml^−1^ streptomycin (Gibco). Cells cultured less than 20 passage numbers were used in the experiments.

### Plasmids and stable cells

FRET biosensor containing vectors: pTriEx4-Rac1-2G (Addgene plasmid # 66110 ; http://n2t.net/addgene:66110 ; RRID:Addgene_66110) and pTriExRhoA2G (Addgene plasmid # 40176 ; http://n2t.net/addgene:40176 ; RRID:Addgene_40176) were both gifts from Olivier Pertz.

Stable cell lines expressing the biosensor constructs were generated as follows. HT1080 cells were co-transfected with the Rac1-2G or RhoA-2G plasmids and the pGL4.21 vector containing a Puro^R^ cassette, using Lipofectamine 2000 (Invitrogen) according to the manufacturer’s instructions. 48 h after transfection, cells were subjected to selection with puromycin (2 μg/ml, Sigma) and subsequently FACS sorted (FACS ARIA, BD) to obtain stably middle-level-expressing cells.

### Polyacrylamide (PAA) gel preparation

PAA gel preparation on glass surface was adapted from previously published protocols^30^ to enable simultaneous imaging of FRET biosensor and red fluorescent beads from the bottom (Fig. 1a). The details of the protocol for polyacrylamide gel preparation including all modifications from previous publications are provided below. 0.1 M NaOH was added to the glass bottom of 35 mm MatTek dish for 5min. The liquid was then removed and the dish was air-dried. ~150 μL 3-aminopropyltrimethoxylsilane (Sigma) was added onto the NaOH treated glass bottom from the previous step for 5 min. The dish was then washed thoroughly with ddH_2_O. Then 2mL 0.5% glutaraldehyde in PBS was added to the MatTek dish for 30 min. The dish was subsequently thoroughly washed with ddH_2_O and air-dried. Another glass coverslip was sequentially coated with 0.1 mg mL^−1^ poly-D-lysine and 1:1000 ddH_2_O diluted red fluorescent beads (F8801, Invitrogen) for 30 min. 6 μl of an acrylamide / bis-acrylamide mixture dissolved in water (the concentrations of crosslinker and polymer were adjusted for a Young’s modulus of 6.9 kPa^29^), 5.6 μg mL^−1^ N-succinimidyl ester of acrylamidohexanoic acid (N6 crosslinker)^38^, 0.5% of ammonium persulphate and 0.05% TEMED was added to the center of the glass bottom of the previously treated MatTek dish. This gel mixture was covered with the red fluorescent beads coated glass coverslip. To ensure that all red beads laid in the top plane of the gel, the dish was flipped during gelation. Once the polymerization was completed, the coverslip was removed carefully. Then the gel was washed twice with PBS and coated with 2 ml of 0.2 mg ml^−1^ rat tail collagen type I (Millipore) at 4 °C overnight.

### Confocal live cell imaging

The collagen type I coated PAA gel was washed with DMEM before usage. Cells were trypsinized, PBS-washed and replated in DMEM (phenol red free) + 1% FBS onto the gel in MatTek dish (4000 cells in the inner well). Cells were then incubated at 37 °C with 5% CO_2_ for 2 h before imaging. Live-cell imaging was performed using a Nikon A1R confocal equipped with GaSaP detectors and environmental chamber. Images were acquired every 10 s for 15 – 40 min, using a 60 × Plan Apo oil objective (1.4 NA) and 1024 × 1024 resolution. FRET biosensor signal acquisition: ‘mTFP1’-channel (457 nm laser excitation, 482/35 nm emission); ‘FRET’-channel (457 nm laser excitation, 525/50 nm emission). Red fluorescent beads signal acquisition: 561 nm laser excitation, 595/50 nm emission. After the time lapse imaging, cells were trypsinized and washed away. Then the fluorescent beads in the same positions were imaged to capture the released state of the PAA gel for TFM reference.

### Lattice light sheet microscopy (LLSM)

The preparation of PAA gel for LLSM was modified from the gel preparation on MatTek dish. The 5 mm round glass coverslip (Warner Instruments, Catalog # CS-5R) was surface activated for gel conjugation and the MatTek glass surface was coated with red fluorescent beads. In this way, the red fluorescent beads labeled PAA gel was conjugated onto the 5 mm diameter coverslip to fit for the LLSM imaging.

The lattice light sheet microscope (LLSM) used in these experiments is housed in the Advanced Imaged Center (AIC) at the Howard Hughes Medical Institute Janelia Research Campus. The system is configured and operated as previously described^39^. Briefly, HT1080 cells were transiently transfected with Rac1 or RhoA biosensor and Lifeact-RFP670 plasmids 16–24 h before imaging using Lipofactamine 3000 (Invitrogen). Cells used for live cell imaging were seeded on the collagen I coated PAA gel conjugated on 5 mm round glass coverslip in Leibovitz L15 Medium, no phenol red (Thermo Fisher Scientific), containing 1% FBS. During imaging, cells were maintained at ~5% CO_2_ concentration, 95% humidity, and 37°C via custom-built environmental chamber (Oko-Labs). Samples were illuminated by a 2D optical lattice generated by a spatial light modulator (SLM, Fourth Dimension Displays). The light sheet pattern was a square lattice with minimum NA of 0.44 and a maximum NA of 0.55. The sample is illuminated by 445 nm, 560 nm, and 642 nm diode lasers (MPB Communications) at 100%, 10%, and 90% AOTF transmittance and 140 mW, 50 mW, and 100 mW initial box power through an excitation objective (Special Optics, 0.65 NA, 3.74-mm WD). Fluorescent emission was collected by detection objective (Nikon, CFI Apo LWD 25XW, 1.1 NA), and a sCMOS camera (Hamamatsu Orca Flash 4.0 v2) at 75 ms exposure time. Acquired data were deskewed as previously described^39^ and deconvolved using an iterative Richardson-Lucy algorithm. Point-spread functions for deconvolution were experimentally measured using 200nm tetraspeck beads adhered to 5 mm glass coverslips (Invitrogen, Catalog # T7280) for each excitation wavelength.

### FRET ratio calculation

FRET image calculation was performed with FIJI^40^. Images from mTFP1 channel and FRET channel were background subtracted and smoothed with median filter. Then the FRET/mTFP1 ratios were calculated and smoothed with median filter to get the final images. 2D median filter and 3D median filter^41^ were used for confocal images and LLSM images respectively.

### Traction force calculation

The traction force calculation was performed with the Matlab based software previously published by the Danuser lab^31^. Briefly, the fluorescent bead channel time-lapse images were registered towards the reference image to correct the stage drift. And then in each time-lapse image, the displacement of the beads towards the reference image was calculated, based on which the traction force was calculated with the Fourier transform traction cytometry method^42^.

### Analysis of image series

The window sampling process was performed with the Matlab based software previously published by the Danuser lab^5^. Briefly, the automatic cell segmentation was performed based on the images of the ‘FRET’ channel. Then the cell edge was sampled into 1 μm wide sectors (or 5 μm for control purpose (Supplementary Fig. 6)). For each sector, 1 μm deep probing windows were created continuously from the cell edge to the cell center based on its local geometry. The local edge sector velocity was obtained, together with the mean Rac1-GTP or RhoA-GTP FRET signals and traction force level in its corresponding windows.

The quality of temporal alignment of the window sampled time lapse images were further improved by comparing the traction force signals in the fifth sampling windows from neighboring time points in each cell. Then the Rac1-GTP/RhoA-GTP and traction force signals from each window were smoothed over time with the Matlab function *csaps* to suppress noise (Fig. 1c; Supplementary Fig. 4). Each time series of the mean GTPase activity and traction force levels obtained from a window provided one instantiation of the dynamics of Rac1-GTP or RhoA-GTP and traction force levels related to the corresponding cell edge sector motion. The alignment of the Rac1-GTP/RhoA-GTP and traction force signals according to the corresponding edge sector protrusion/retraction onset/V_max_ was performed by following the previously published methods^5^. Briefly, the protrusion and retraction time series of the edge sector were acquired based on the local maxima/minima of the edge sector displacement. Protrusions/retractions with short distances (<1 μm) or time periods (<1 min) were discarded. Remaining protrusion/retraction time series were aligned according to the edge protrusion/retraction onset/V_max_ events and the mean values and 95% confidence intervals of the corresponding Rac1-GTP/RhoA-GTP and traction force around the four different events were calculated.

We focused most of our analysis on the windows located 1-2 μm from the segmented cell edge because we found them to most likely represent the actual cell edge (due to segmentation imperfection) and to entail the most meaningful dynamics of Rac1-GTP, RhoA-GTP and traction forces (see methods; Fig. 1d; and Supplementary Fig. 6).

Although the absolute value is different, we found that the traction force in the second depth layer sampling windows were positively correlated to the traction force 4-6 μm away from cell edge with no time delay (Supplementary Fig. 6). Therefore, we also used signals in the second sampling windows for further analysis of traction force change dynamics, thereby also measuring and relating the RhoA-GTP, or Rac1-GTP and traction forces in the exact same locations.

### Optogenetic perturbation

HT1080 cells stably expressing mCherry-tagged photoactivatable constitutively active (GTP-bound) Rac1 (PA-Rac1(Q61L)) or dominant negative (GDP-bound) Rac1 (PA-Rac1(T17N))^34^ were seeded onto the collagen type I coated PAA gel surface-labeled with far-red fluorescent beads. 2 h after cell seeding, local Rac1-GTP activity of a small region of the cells were elevated (PA-Rac1(Q61L)) for 1 min or inhibited (PA-Rac1(T17N)) for 2 min with pulsed 488 nm laser. Cell membrane dynamics (mCherry) and traction force (0.2 μm far-red beads, Invitrogen) images were obtained using an environmental chamber equipped confocal microscope imaging at 5 s intervals from 5 min before the perturbation started until 5 min after the perturbation finished. After the time-lapse imaging, cells were detached from the PAA gel with trypsin and reference images in the far-red channel were acquired for traction force calculation.

### Difference in difference (DID) method

The mean protrusion or retraction velocity/distance of the membrane edge sector time series in the perturbation region before photoactivation were extracted and aligned according to the onset time points as unperturbed controls. The mean time from protrusion or retraction onset until V_max_ reaching were obtained from these control protrusion or retraction velocity curves. The edge sectors which started to protrude or retract close to the perturbation starting time point were extracted and grouped according to the differences of their perturbation starting time and mean V_max_ reaching time. The mean velocities/distances of the perturbed protrusion or retraction time series in each group were calculated and their differences towards corresponding control time series were then plotted.

### Quantification of traction force changes after optogenetic perturbation

To compare the traction force change with and without optogenetic perturbation in the photoactivated cell region, mean traction forces of each cell in the perturbed region before (0-1 min) and after (3-4 min) the perturbation starting time point were quantified. For each cell, the mean traction force after perturbation was normalized to the mean traction force before perturbation. Then the normalized traction force changes after perturbation were compared after PA-Rac1(Q61L) or PA-Rac1(T17N) induction.

To compare the traction force change after optogenetic perturbation in protrusion and retraction windows, cells were masked, and the windows were sampled according to the signal from the mCherry channel. Protrusion and retraction sectors starting near the perturbation starting time point were selected and aligned in the same way as described above. Control protrusion and retraction sectors were obtained based on the time lapse images before the perturbation started. Then the mean traction force change 2 min after the start of induction of PA-Rac1(Q61L) or PA-Rac1(T17N) in the second window (1-2 μm from the segmented cell edge) of the corresponding protrusion or retraction sectors initiated within 20 s before the perturbation started were compared to the mean traction force change in the windows of the sectors before the perturbation started, at time points when these sectors displayed corresponding phases of protrusions or retractions.

### Statistics

The 95% bootstrap confidence intervals of the mean value of the aligned time series were calculated with bootstrap resampling method^43^ (Matlab function *bootci*).

The false discovery rate (FDR) method^44, 45^ was performed with own Matlab scripts. Because of the large difference of sample size between control (>1000) and perturbed (~50-100) groups, the same number of time series from the control group was randomly sampled towards the perturbed group for 10000 times. For each time, the standard deviations of the grouped time series (control and perturbed) were compared to the standard deviation of entire two-group time series with two sample t-test for a *p* value at each time point. And the grouped data were also randomly regrouped and standard deviation compared for a control *p* value at each time point. After the 10000 time comparations, the Q values of positive FDR threshold was calculated (Matlab function *mafdr*()) based on the *p* values derived from both grouped data and randomized data. The 0.1% threshold of Q value was defined from the randomized data and the mean Q value from the grouped data was compared to the threshold for significance.

The paired *t*-test was performed with Matlab (function *ttest*()) and the two-sample *t*-test was performed with Matlab (function *ttest2*()).

## Acknowledgements

This study was supported by The Swedish Foundation for Strategic Research (SB16-0046 (Sysmic)), The Swedish Research Council and The Swedish Cancer Society. Confocal microscopy was performed at the LCI facility/Nikon Center of Excellence, Karolinska Institutet (KI), Sweden supported by grants from the KI infrastructure committee and the Centre for Innovative Medicine at KI. The LLSM imaging data were produced in collaboration with the Advanced Imaging Center, a jointly funded venture of the Gordon and Betty Moore Foundation and the Howard Hughes Medical Institute at HHMI’s Janelia Research Campus. We thank Xavier Serra-Picamal, KI, for experimental advice, Pontus Nordenfelt, Lund University, Sweden, for analysis advice, Wolfgang Huber, EMBL, Germany, for statistic advice and Peter Friedl, Radboud Univeristy, The Netherlands and Onur Dagliyan, Harvard Medical School, USA for helpful comments on the manuscript. Schematic images (Figure 1a and Figure 4) were created with BioRender.com.

## Data availability

All data supporting the findings of this study are available from the corresponding author upon reasonable request.

## Code availability

Custom codes used in this study have been deposited on GitHub (https://github.com/hujianjiang/Rac1).

## Author Contributions

J.H. conceived, designed, and performed most experiments and analyses. J.H. performed confocal live cell imaging and related image analyses. J.H. and X.G. performed LLSM live cell imaging. J.H. performed the LLSM image analysis. J.H. performed image analyses, statistical analyses and data visualization. J.H. and S.S. wrote the manuscript and all authors approved the final version.

## Supplementary information

**Supplementary Figure 1.**
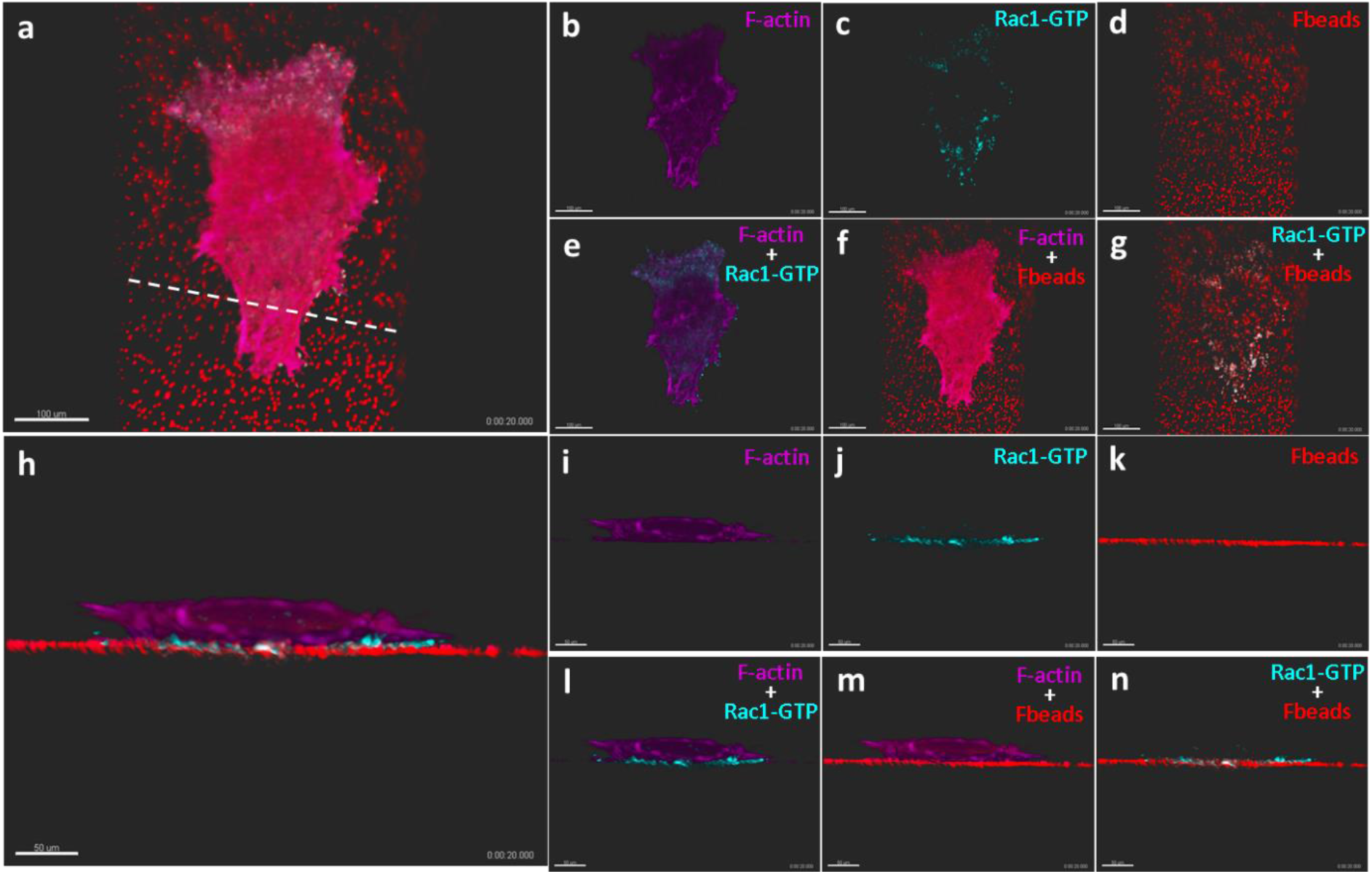
Lattice light sheet microscopy displays Rac1-GTP at the cell-matrix interface. HT1080 cells expressing a Rac1-GTP biosensor and miRFP670 tagged F-actin were grown on a Collagen type I coated PAA gel surface labeled with red fluorescent beads. Rac1-GTP levels (cyan), fluorescent beads (red) and F-actin (purple) were imaged in 3D in live cells by lattice light sheet microscopy. Upper images (**a-g**): xy view of a sample cell, scale bar 100 μm. Lower images (**h-n**): side view of the sample cell via the white dash line in **a**, scale bar 50 μm. Left large images (**a, h**) show the composition of all three channels. Small images on the right show the single channels (**b-d, i-k**) and composed images of two channels at the time (**e-g, l-n**).

**Supplementary Figure 2.**
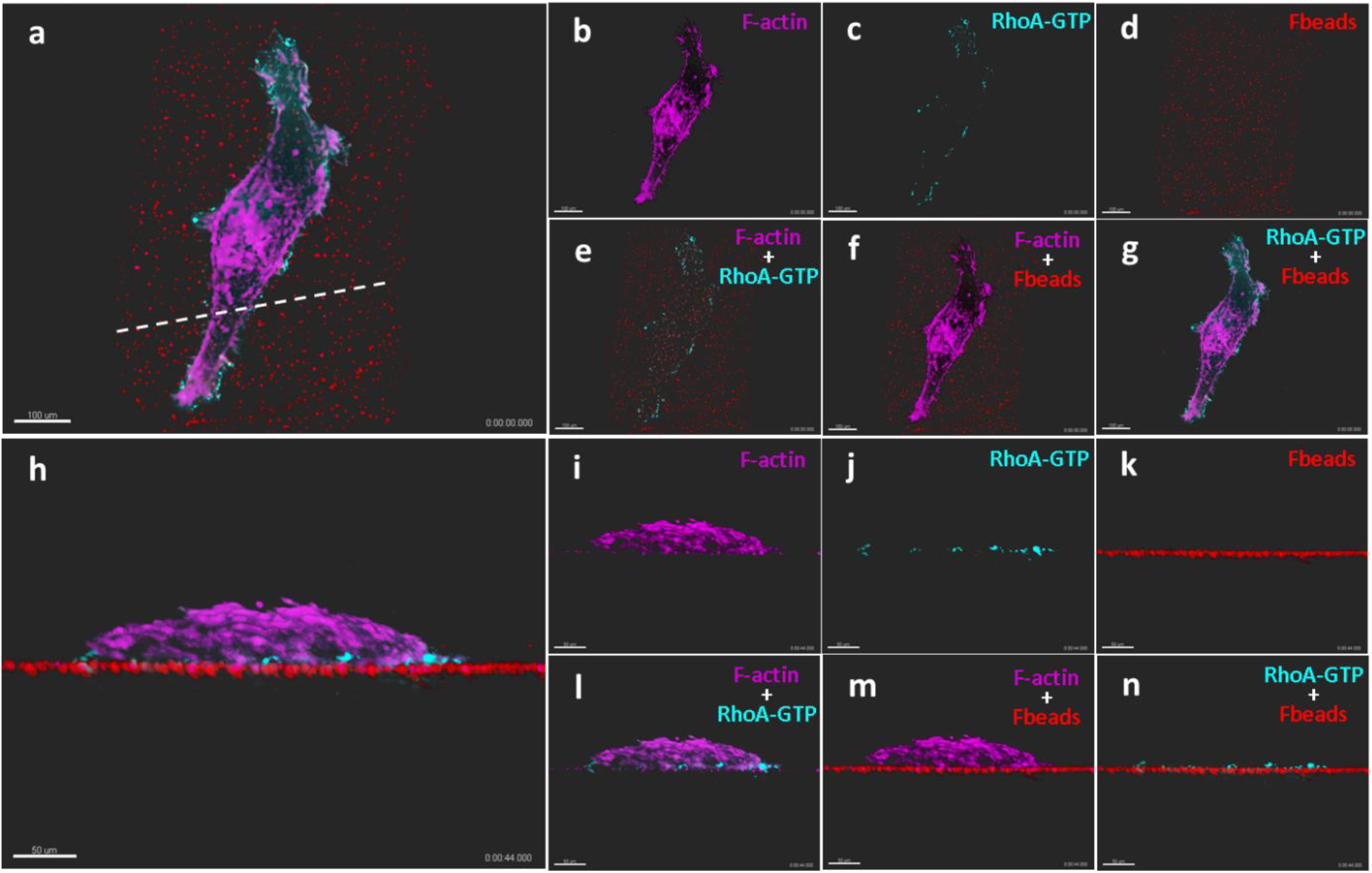
Lattice light sheet microscopy displays RhoA-GTP at the cell-matrix interface. HT1080 cells expressing RhoA biosensor and miRFP670 tagged F-actin were grown on a Collagen type I coated PAA gel surface-labeled with red fluorescent beads. RhoA-GTP levels (cyan), fluorescent beads (red) and F-actin (purple) were imaged in 3D in live cells by lattice light sheet microscopy. Upper images (**a-g**): xy view of a sample cell, scale bar 100 μm. Lower images (**h-n**): side view of the sample cell via the white dash line in **a**, scale bar 50 μm. Left large images (**a, h**) show the composition of all three channels. Small images on the right show the single channels (**b-d, i-k**) and composed images of two channels at the time (**e-g, l-n**).

**Supplementary Figure 3.**
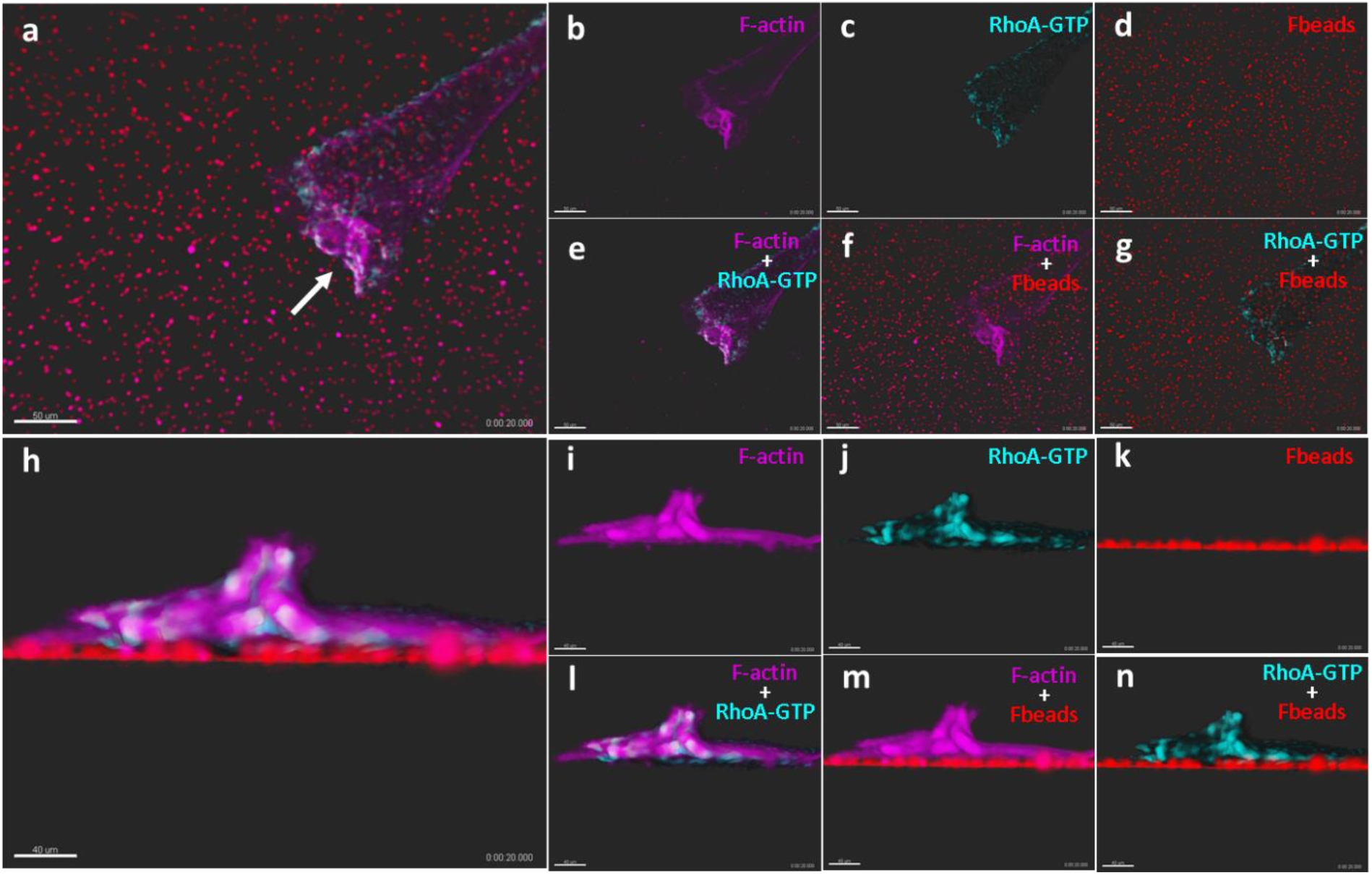
Lattice light sheet microscopy displays RhoA-GTP at membrane ruffles. HT1080 cells expressing RhoA biosensor and miRFP670 tagged F-actin were grown on a Collagen type I coated PAA gel surface-labeled with red fluorescent beads. RhoA-GTP levels (cyan), fluorescent beads (red) and F-actin (purple) were imaged in 3D in live cells. Upper images (**a-g**): xy view of a sample cell with a protrusion membrane ruffle, scale bar 50 μm. Lower images (**h-n**): side view of the sample cell via the white arrow in **a** that points to a protrusion ruffle with high RhoA-GTP levels above the cell-ECM interface, scale bar 40 μm. Left large images (**a, h**) show the composition of all three channels. Small images on the right show the single channels (**b-d, i-k**) and composed images of two channels at the time (**e-g, l-n**).

**Supplementary Figure 4.**
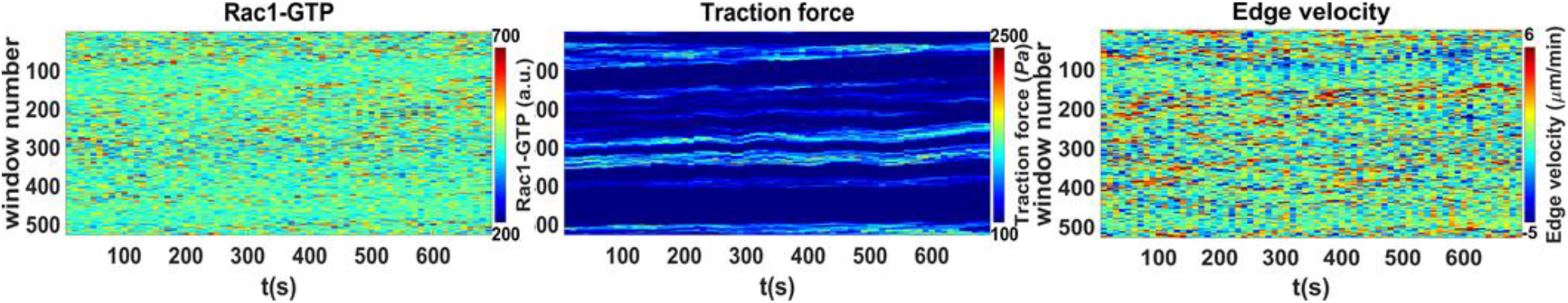
Kymographs of GTPase and traction force levels and the corresponding edge velocity before alignment and smoothing. Temporal dynamics of Rac1-GTP levels (left) and traction force (middle) levels in the second depth windows (1-2 μm from the cell edge) of a cell, as well as corresponding edge velocity (right) of each sector without alignment and smoothing. All graphs display pseudo colors as indicated in the insets. Corresponds to registered and smoothed graphs in Fig 1c.

**Supplementary Figure 5.**
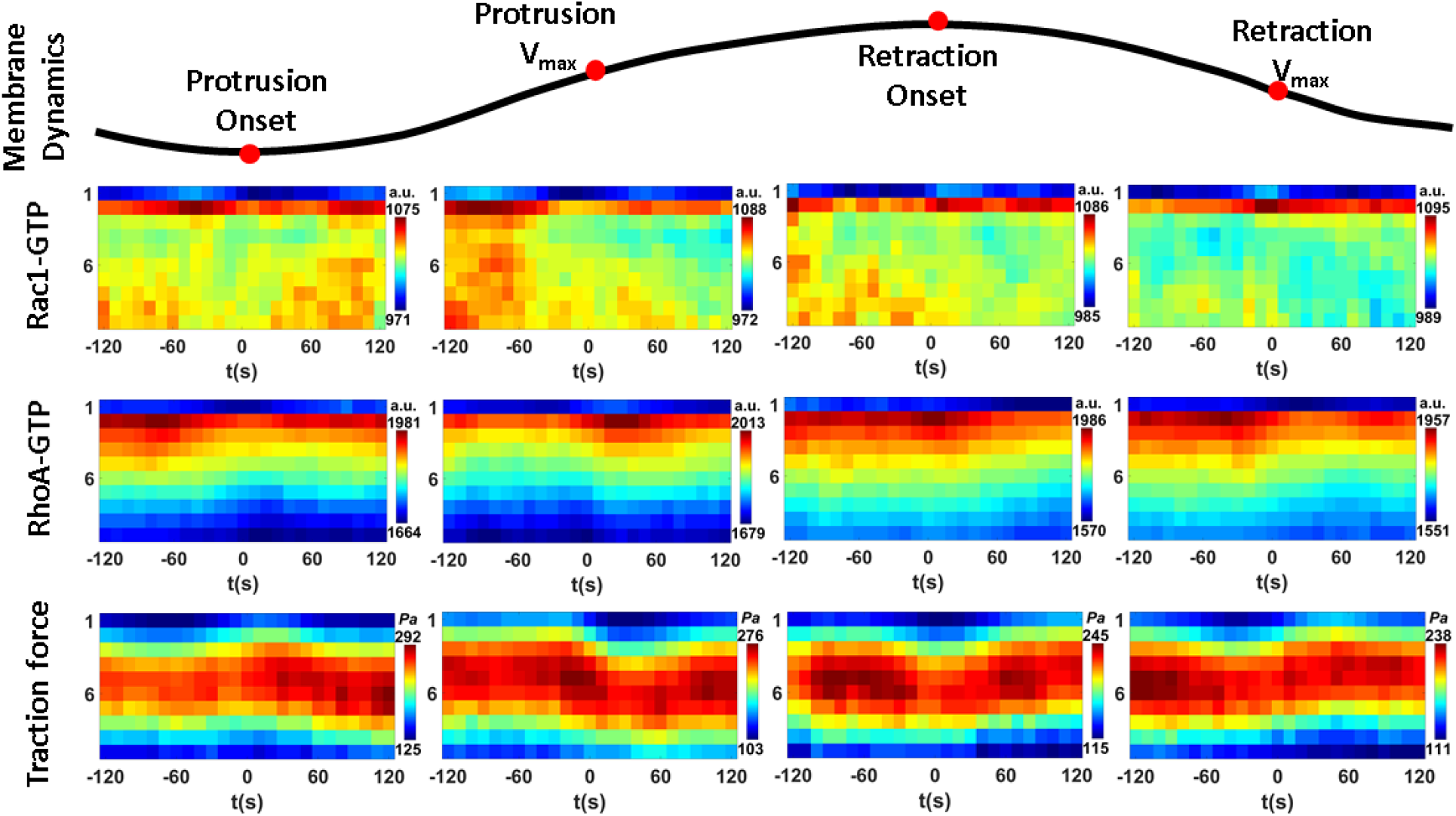
Rac1-GTP, RhoA-GTP and traction force levels around key cell membrane events analyzed with an alternative sample window size. The dynamics of 5 μm wide segmented cell membrane edge sectors were aligned according to cell membrane protrusion onset, maximal protrusion velocity (protrusion V_max_), retraction onset, and maximal retraction velocity (retraction V_max_). The mean values of Rac1-GTP, RhoA-GTP and traction force dynamics at different depths of the cell (1-10 μm, y axis) around the specific time points (−120 s to + 120 s, x axis) are shown in pseudo-color. Sample size: Rac1-GTP & traction force: 1056 protrusions and 796 retractions from 13 cells; RhoA-GTP: 1631 protrusions and 1023 retractions from 23 cells. The outcome is almost identical with that using 1 μm wide sector (Fig. 1d) showing that the results are insensitive to the segmentation sector width up to 5 μm.

**Supplementary Figure 6.**
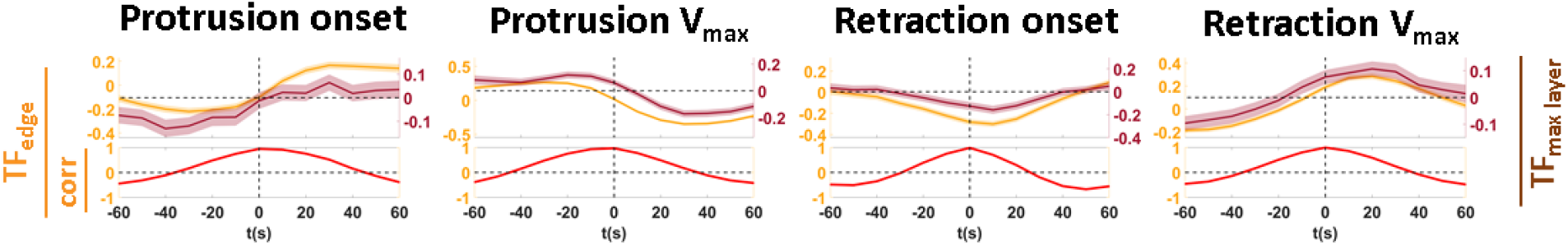
High correlation of traction force dynamics at different cell depth windows. The highest traction force levels occurred 4-6 um away from the cell edge. Here we show cross-correlation analysis of mean traction force levels in the second depth window (TF_edge_, 1-2 μm from the segmented cell edge) with the corresponding mean traction force levels in the 4th and 5th depth layers of the window (TF_max layer_, 4-6 μm away from cell edge) during protrusion onset, protrusion V_max_, retraction onset and retraction V_max_. For the above paired data, Solid lines show the mean values and the shadows show the 95% confidence interval. Correlation results were shown in red lines below the corresponding pairs.

**Supplementary Movie 1.**
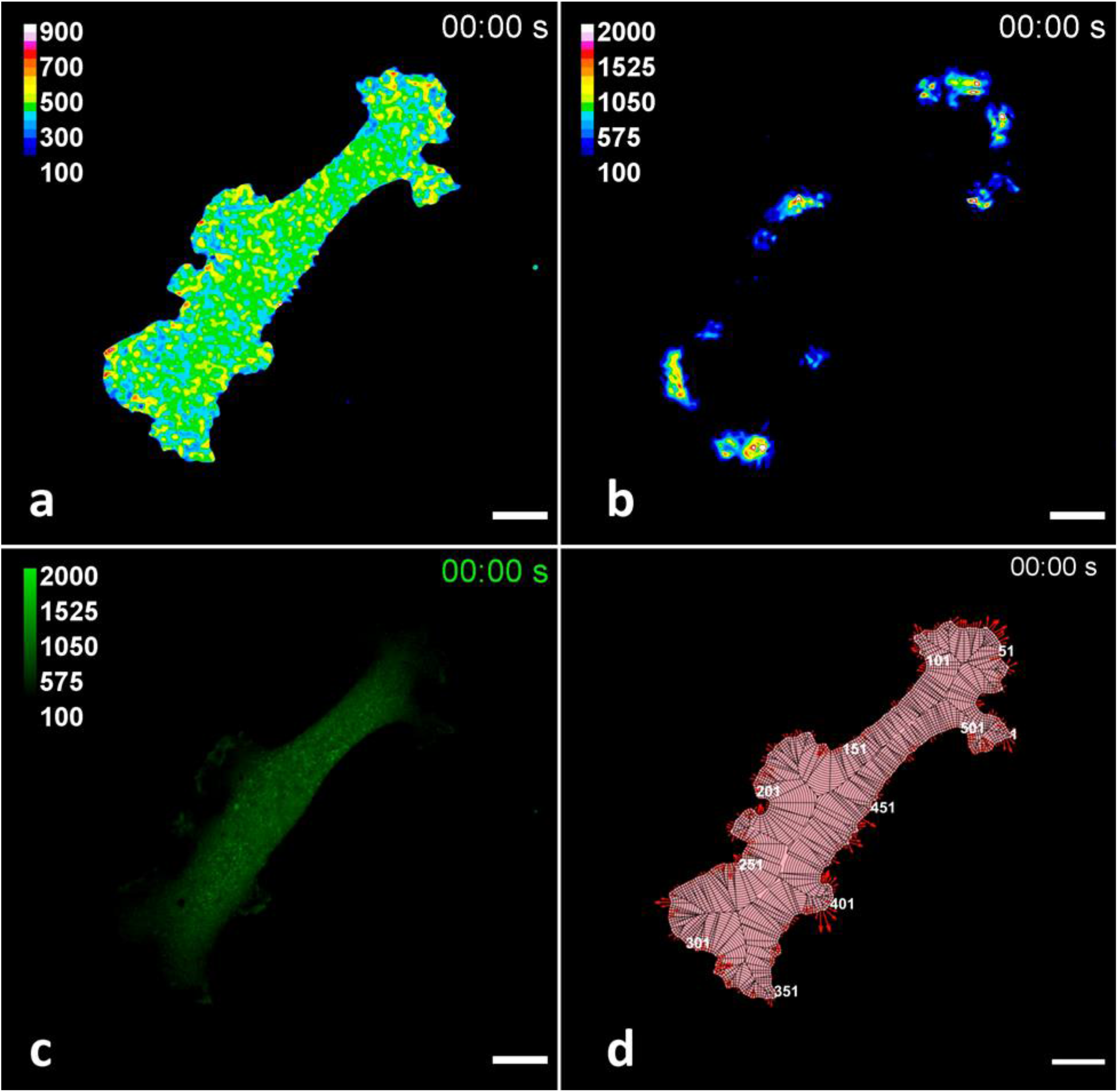
Time lapse movies corresponding to images in Figure 1b. The sample time lapse movies show the measured Rac1-GTP level (a), traction force (b), signals in the yfp channel (c) and outlay of the window sampling results (d) during cell migration. Rac1-GTP (a.u.) and traction force (*Pa*) levels are shown in pseudo colors according to the inset scales. Signals in the yfp channel indicate the distribution of Rac1 biosensor in the cell. For the window sampling images, the pink area shows the segmented cell area and the black lines show how the cells are sampled into 1 μm wide cell edge windows based on local cell edge geometry. Windows are numbered in white along the cell edge around the cell. Red arrows show the instant edge sector velocity. Scale bar 20 μm.

